# MBCO PathNet: Integration and visualization of networks connecting functionally related pathways predicted from transcriptomic and proteomic datasets

**DOI:** 10.1101/2025.02.19.638900

**Authors:** Jens Hansen, Ravi Iyengar

## Abstract

Our desktop application MBCO PathNet allows for quick and easy integration and visualization of networks of functionally related pathways predicted from numerous gene and protein lists using the Molecular Biology of the Cell Ontology (MBCO) and other ontologies. Within networks of hierarchical parent-child relationships or functional relationships, pathways are visualized as pie charts where each slice represents a dataset that predicted that pathway. Sizes of pies and slices can be selected to represent statistical significance or other quantitative measures. In addition, MBCO PathNet can generate bar diagrams, heatmaps and timelines.

**Availability and implementation:** mbc-ontology.org

github.com/SBCNY/Molecular-Biology-ofthe-Cell

iyengarlab.org/mbcpathnet

## Introduction

Whole cell physiological functions are generated by coordinated interactions between pathways that form functional networks. Each pathway comprises multiple gene products that interact with each other to generate a subcellular activity. High-throughput (HT) technologies such as single cell (sc), nucleus (sn) and spatial transcriptomics or regional proteomics have remarkably increased the opportunities to investigate whole cell physiological dynamics. To reveal mechanistic insight that is not easily identified from initial lists of genes, pathway enrichment analysis allows for hundreds of experimentally obtained gene products to be organized as pathways summarizing the overrepresented functions and activities. Using prior knowledge from selected ontologies, e.g. Gene Ontology (GO) ^1,2^ and Reactome,^3^ pathway enrichment analysis identifies pathways that are statistically enriched in genes with altered expression. The hierarchical organization of ontologies in parent-child relationships, where the children describe sub-functions of their parents, allows documentation of taxonomical relationships between predicted pathways for the same and different datasets, simplifying the interpretability of the results. To characterize networks of functionally interacting subcellular processes (SCPs) underlying whole cell functions, we have developed the Molecular Biology of the Cell Ontology (MBCO).^4^ Its annotated hierarchy spans three to four SCP levels where SCPs of the same level describe distinct functions of similar biological details as found in the textbook. Annotated parent child relationships are enriched with interactions between functionally related SCPs that were inferred from prior knowledge. Consideration of these interactions allows a new enrichment approach, dynamic enrichment analysis, which considers interactions and dependencies between the enriched SCPs.

To address the need for quick integration of numerous multi-dimensional datasets such as sc transcriptomic datasets using MBCO and other ontologies, we have developed a desktop application MBCO PathNet. Gene and protein lists obtained by various software packages analyzing HT experiments can be easily uploaded and subjected to pathway identification. The application allows both standard and dynamic enrichment analysis using MBCO, GO, Reactome and two user-supplied ontologies. Multiple datasets can be quickly combined into groups of interest to enable their integrative analysis within networks of pathways. Results can be shown as bardiagrams, timelines or heatmaps. Genes that show experimentally measured changes can be mapped to identified pathways and added to the networks as child nodes. The application also allows focusing on selected SCPs or grouping them into new higher-level SCPs that underlie whole cell functions.

## Implementation

Within windows the application can be started by opening the exe-file, within a LINUX environment it can be opened using Mono (mono-project.com), a software platform for the Microsoft NET environment that was sponsored by Microsoft. See github link for related terminal commands.

A central element within our application is the quick assignment of datasets, i.e. user-supplied lists of genes or proteins, to dataset integration groups. Integration groups should contain those datasets that the user wants to cross-compare. Enrichment results obtained for datasets within the same integration group will be visualized in the same figures. SCP networks predicted for datasets of the same integration group will be merged and cross-connected. Integration groups can either be defined using the functionalities of the application within the “Organize data” menu (see below) or be added to the user-supplied data before import.

User-generated datasets can be uploaded into the application, using two different ways. Gene or protein lists can either be copy pasted into the Gene list box or be read automatically from text files, using the “Read data” menu. The “Read data” menu allows quick upload of tens to hundreds of different datasets into the application. Depending on the user’s selection the application will either read a single file or all files within a given directory. The eight text boxes in the menu panel allow specification of column names in user-supplied files that map to selected fields within our application, e.g. the integration group field. Fields with empty text boxes will be ignored during the upload. The application will search each supplied file for specified column names and return an error message, if a name was not found. Files whose names end with “_bgGenes” are assumed to contain a single column of genes or proteins without a headline that will be imported as experimental background genes. The experimental background genes will be automatically mapped to all datasets uploaded from a file with the same name, except the “_bgGenes” ending. Experimental background genes should contain all genes or proteins that were tested for significance, e.g. for being differentially expressed. In an RNA sequencing experiment, a potential selection for background genes could be all genes that are annotated to the reference genome, but more strict definitions are also reasonable. Their inclusion improves statistical accuracy. Upload and mapping of background genes can additionally be achieved using the menu “Background genes”. To allow quick reproduction and recapitulation of results from previously analyzed datasets, analyzed gene/protein lists, associated sets of background genes and user selected parameter settings can be automatically re-uploaded into the application. Every time a dataset is submitted to analysis, the application will write related files into the result subdirectory “Input_data”. Automatic reimport is enabled after specification of that directory as the data source and selection of the “MBCO” default column names.

The next menu, “Set data cutoffs”, allows for definition of significance cutoffs that determine which genes of each dataset will be subjected to enrichment analysis.

One of the main functions offered by the “Organize data” menu is the quick assignment of integration groups and colors to the user-supplied datasets as well as their deletion. This is enabled by temporal grouping of datasets based on selectable shared characteristics. Described actions will be applied to all datasets within the same temporal group. Integration groups and colors can automatically be assigned. Among the selectable characteristics are shared time points, directionalities of change (e.g., up- or downregulation) or shared user-selected substrings within the dataset names. Based on a user-supplied delimiter, the application will internally split the dataset names into multiple substrings. For example the name “LMD RNASeq - CD – IC” will be split into “LMD RNASeq”, “CD” and “IC”, if “– “was selected as a delimiter. The user can then specify the position index of the relevant substring, counted from the left and/or the right. For definition of multiple substrings whose combined matching will define a temporal group the user can separate the indices by commas. As another functionality, the menu “Organize data” allows the specification of the order of the results for datasets of the same integration group in bar diagrams, heatmaps and pie charts.

The menu “Enrichment” allows specification of enrichment parameters and selection of types of figures to be generated for display of results. User-supplied datasets are subjected to standard enrichment analysis that is based on the right-tailed Fisher’s exact test. This menu allows selection of the ontology to be used as well as definition of two SCP significance cutoffs, i.e., maximum significance p-value and rank. Pathways are significant, if they meet both cutoffs. Since MBCO SCPs are in parent and child relationships across the different levels but show minimal redundancies between SCPs within the same level, the application enables the definition of cutoffs for each SCP level separately. In addition, MBCO allows dynamic enrichment analysis. Briefly, dynamic enrichment analysis merges up to three functionally related SCPs that contain at least one gene of the analyzed dataset into dataset-selective higher-level SCPs and adds those to a dataset-selective ontology, before enrichment analysis using Fisher’s exact test. SCP cutoffs for dynamic enrichment analysis can be separately specified. Functional SCP relationships are defined by the weighted inferred MBCO SCP interactions and the user can select the percentage of the top interactions to be considered. In the case of the three GO namespaces, i.e., biological processes, cellular components and molecular functions, the “Enrichment” menu allows removal of too large or too small GO terms by specification of a maximum and minimum number of genes within a considered GO term.

The functionalities of the menu “SCP-networks” allow to select, if predicted SCPs of the same level shall be boxed and if SCPs should be connected across all datasets of the same integration group based on parent-child and/or functional relationships. Percentages of considered top inferred MBCO interactions can be specified for standard enrichment analysis, while they are the same as those specified in the menu “Enrichment” for dynamic enrichment analysis. Genes with altered expression that map to predicted pathways can be added to the networks as child nodes. Since the hierarchical organization of other ontologies often contains parent-child relationships over multiple levels, the user can specify, if predicted pathways shall only be connected by intermediate nodes or if all of their ancestors should be added. In the case of GO, only ancestors are considered that fulfill the size requirements specified in the menu “Enrichment”. Additionally, GO pathways can additionally be connected based on regulatory interactions. SCP nodes will be visualized as pie charts where each slice represents one dataset that predicted that SCP meeting the significance criteria specified in the menu “Enrichment”. The user can select the slice and pie chart areas to be proportional to each −log_10_(p-value) and the sum of all −log_10_(p-values) predicted for related datasets, respectively. −log_10_(p-values) predicted by dynamic enrichment analysis will be split equally among all contributing SCPs and multiple split −log_10_(p-values) for the same SCP will be summed up. For further emphasis on SCPs predicted with high significance and by many datasets, the application allows selecting that SCP label sizes increase with increasing SCP node areas. Label size ranges can be specified. As alternative approaches for visualization of results in pie charts, pie chart areas can be selected to be proportional to the total number of related datasets or the number of related datasets assigned to the same color. Same color assignments can for example be used to highlight the HT assays, if each generated multiple datasets (e.g., to compare sc and sn transcriptomics, if each assay predicted multiple subtypes for a cell type of interest). As a last option, the user can select that pie charts have identical sizes for all SCPs. In case of the three last discussed selections slice sizes will be identical within the same pie chart. Node scaling can be set to be unique for each network or to be the same for all networks predicted by standard or dynamic enrichment analysis.

The menu “Select SCPs” allows grouping of SCPs or hierarchical SCP branches that will be shown in all result figures and networks, if the check box at the bottom of the menu is activated. Results figures or networks will be generated for all SCPs of the same group, while significance cutoffs will be ignored. To analyze which dataset genes map to selected SCPs, dataset genes can be added as child nodes to the SCPs.

Within the menu “Define own SCPs” the user can generate new SCPs by merging existing SCPs, which will be add to the ontology.

Networks generated by MBCO PathNet can be visualized with yED graph editor.^5^

## Methods

After starting the application, we opened the “Example data” menu and loaded the “LINCS DtoxS / predicTox examples” datasets. We opened the “Enrichment” menu and selected “GO biological process” in the “Ontology” list box at the top of the application. As instructed by the application, we downloaded the “go-basic.obo” and “goa_human.gaf” files from www.geneontology.org (Feb 1, 2025) and copied both files into the “Other_datasets” folder. Selecting “GO biological process” again instructed the application to populate all GO parent terms with their child term genes and save the results for reuse. We left all parameters as updated by the application when loading the example data. Pressing the “Analyze” button generated and saved a populated GO biological process annotation file that meets the minimum and maximum size requirements specified in the “Enrichment” menu and started the analysis. As instructed by the application, we opened the “Results/SCP_networks/” folder and opened the “Go_bp_human_prednisolone - Down_standard.graphml” file with yED, ^5^ selected “Tree” ‘– “Directed” in the “Layout” menu and activated “Consider Node Labels” in the “Directed” tab in the appearing pop-up window. SCP nodes were rearranged manually. Finally, we selected “Edge routing” – “Orthogonal/Polyline” in the “Layout” menu. Next, we selected “MBCO human” in the “Ontology” list box of the “Enrichment” menu of our application and deactivated “Label sizes ∼ Node sizes” in the “SCP networks” menu. Pressing the “Analyze” button started the analysis. Generated SCP networks for standard (and dynamic) enrichment analysis of bortezomib-upregulated genes were processed using yED, as described above. Within the “SCP networks” menu of our application, we now reactivated “Label sizes ∼ Node sizes” and pressed the “Analyze” button for a third time. SCP networks for dynamic enrichment analysis of lapatinib-upregulated genes were prepared as described above.

## Results

We have previously used the functionalities now implemented in MBCO PathNet to integrate multiomics datasets obtained from undiseased human reference kidney samples within the Kidney Precision Medicine Project (KPMP),^6^ for the documentation of SCP networks triggering neurite outgrowth ^7^ and the description of potential mechanisms associated with cardiotoxic tyrosine kinase inhibitors ^8^ within the LINCS program. Using the cardiotoxicity study, we reanalyzed the gene expression profiles ^8,9^ obtained for prednisolone, bortezomib and lapatinib in six healthy human subject iPSC-derived cardiomyocytes. Enrichment analysis of genes downregulated by prednisolone using GO Biological Processes aligns with prednisolone’s inhibitory effect on cell cycle progression ^10^ (Fig. 1A). Our application projected the top five predicted GO Biological Processes with a minimum and maximum number of five and 350 annotated genes into the GO directed acyclic graph to document parent-child relationships. Excluding the top processes predicted for the cell line MSN02 (not shown), all processes describe mitotic chromosome and nuclear division events, aligning with prednisolone’s anti-proliferative effect. Genes upregulated by the proteasome inhibitor bortezomib were re-submitted to standard enrichment analysis using MBCO. Building on our first analysis, ^8^ we here integrated the top five, five, ten and five level-1, -2, -3, and -4 SCPs into the MBCO hierarchy of parent-child relationships. Many of the SCPs predicted with high significance mapped to the “Intracellular degradation pathways” branch, suggesting a compensatory response to overcome proteasomal inhibition (Fig. 1B). Level-3 SCP networks generated by dynamic enrichment analysis (using the top five predictions) contained almost exclusively SCPs related to proteasomal degradation and autophagy (not shown), further supporting this hypothesis. Genes upregulated by lapatinib were subjected to dynamic enrichment analysis using MBCO. Top five level-3 predictions identify lapatinib’s influence on cholesterol homeostasis to an even greater extent than our initial results obtained by standard enrichment analysis.^8^ In agreement, lapatinib-resistance has been linked to upregulated mevalonate pathway activities in cultured breast cancer cells that could be reversed by statin treatment.^11^

**Fig. 1.**
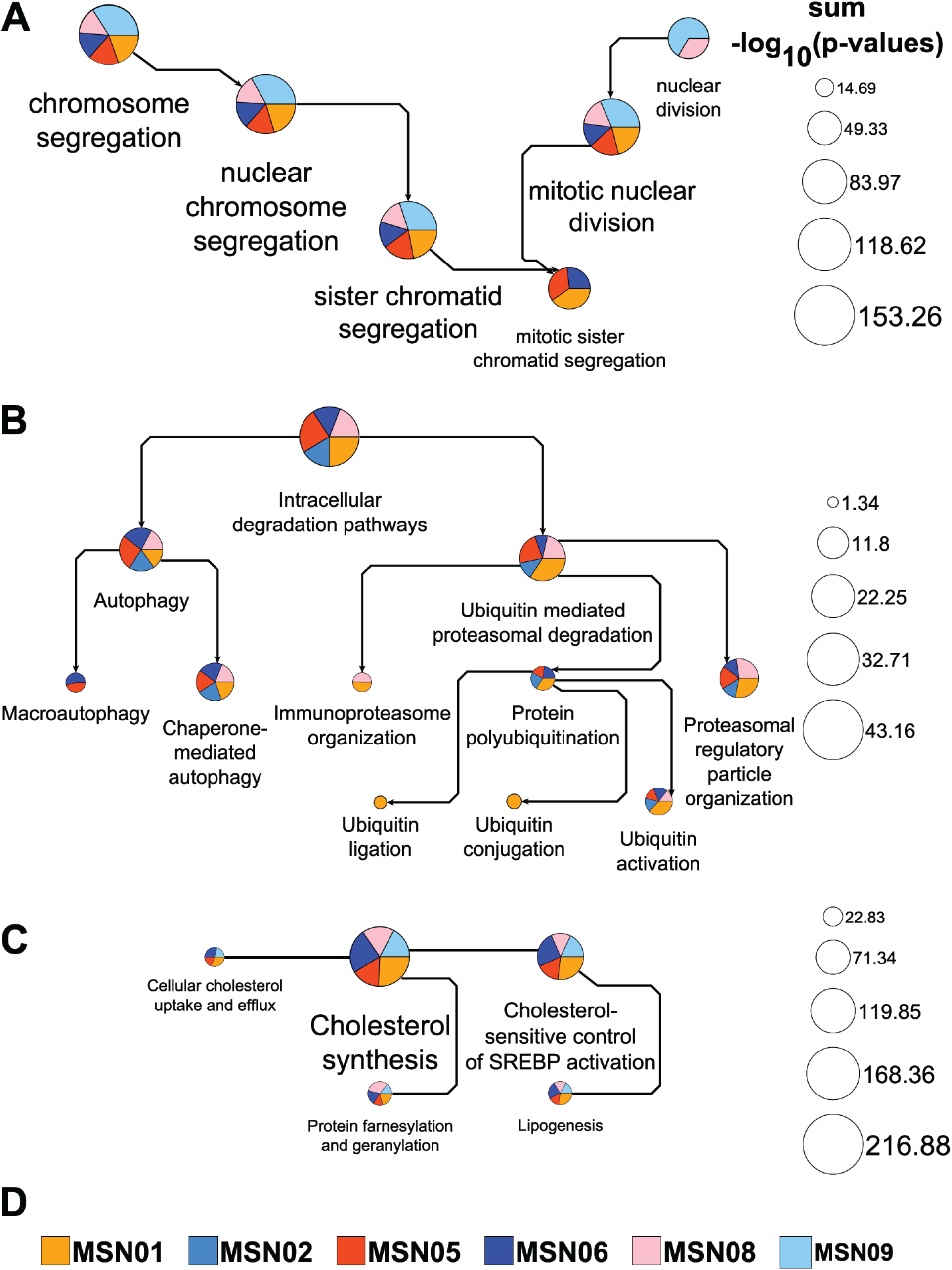
Pathway networks generated from drug-induced up- or downregulated genes. Up- or downregulated genes among the top 600 differentially expressed genes induced by three example drugs in up to six healthy human subject iPSC-derived cardiomyocyte cell lines were subjected to pathway enrichment analysis using MBCO PathNet. **(A)** The top five downregulated GO Biological Processes (min/max # pathway genes: 5/350) in five out of six treated cell lines were integrated into the GO directed acyclic graph. Results obtained for the sixth cell line describe extracellular matrix processes and are excluded from the figure. Pie chart slice and total areas are proportional to individual and the sum of all shown -log_10_(p-values), respectively. Arrows point from parent to child processes. Process labels were selected to increase and decrease with pie chart sizes. **(B)** The top five, five, ten and five level-1, -2, -3 and -4 MBCO SCPs upregulated by bortezomib in five cell lines were integrated into the MBCO parent-child hierarchy. Enrichment results are shown for the SCP “Intracellular degradation pathways” and all its descendants. Each arrow points from a parent to its child SCP, allowing identification of SCP levels by counting the arrows that separate an SCP from the overall level-1 parent. Note that we selected unique SCP label sizes for this visualization. **(C)** Enrichment analysis of lapatinib-upregulated genes in five cell lines using MBCO and dynamic enrichment analysis predicts a level-3 SCP-network focusing almost exclusively on cholesterol-related SCPs (generated from the top five predictions). SCPs are connected, if they belong to the top 25% inferred interactions between level-3 SCPs that were also used as the network basis for dynamic enrichment analysis.

In summary, MBCO PathNet allows quick and easy upload, analysis and integration of multiple omics datasets. Generated SCP networks help with fast and simultaneous investigation and interpretation of the enrichment results for multiple datasets to identify the cell level processes that are affected by the experimental condition.

## Acknowledgements

This project was supported by NIH grant R01-GM137065. We thank Pedro Martinez for the design and set up of the webpage for MBC PathNet.

## References

1 Gene Ontology, C. et al. The Gene Ontology knowledgebase in 2023. Genetics 224 (2023). 10.1093/genetics/iyad031

2 Ashburner, M. et al. Gene ontology: tool for the unification of biology. The Gene Ontology Consortium. Nat Genet 25, 25–29 (2000). 10.1038/75556

3 Milacic, M. et al. The Reactome Pathway Knowledgebase 2024. Nucleic Acids Res 52, D672–D678 (2024). 10.1093/nar/gkad1025

4 Hansen, J., Meretzky, D., Woldesenbet, S., Stolovitzky, G. & Iyengar, R. A flexible ontology for inference of emergent whole cell function from relationships between subcellular processes. Sci Rep 7, 17689 (2017). 10.1038/s41598-017-16627-4

5 yWorks GmbH. yED graph editor, www.yworks.com. Tübingen, Germany

6 Hansen, J. et al. A reference tissue atlas for the human kidney. Sci Adv 8, eabn4965 (2022). 10.1126/sciadv.abn4965

7 Hansen, J. et al. Whole cell response to receptor stimulation involves many deep and distributed subcellular biochemical processes. J Biol Chem 298, 102325 (2022). 10.1016/j.jbc.2022.102325

8 Hansen, J. et al. Multiscale mapping of transcriptomic signatures for cardiotoxic drugs. Nat Commun 15, 7968 (2024). 10.1038/s41467-024-52145-4

9 predicTox.org.

10 Mattern, J., Buchler, M. W. & Herr, I. Cell cycle arrest by glucocorticoids may protect normal tissue and solid tumors from cancer therapy. Cancer Biol Ther 6, 1345–1354 (2007). 10.4161/cbt.6.9.4765

11 Sethunath, V. et al. Targeting the Mevalonate Pathway to Overcome Acquired Anti-HER2 Treatment Resistance in Breast Cancer. Mol Cancer Res 17, 2318–2330 (2019). 10.1158/1541-7786.MCR-19-0756

